# An isogenic panel of single *App* knock-in mouse models of Alzheimer’s disease confers differential profiles of β-secretase inhibition and endosomal abnormalities

**DOI:** 10.1101/2021.08.22.457278

**Authors:** Naoto Watamura, Kaori Sato, Gen Shiihashi, Ayami Iwasaki, Naoko Kamano, Mika Takahashi, Misaki Sekiguchi, Naomi Yamazaki, Ryo Fujioka, Kenichi Nagata, Shoko Hashimoto, Takashi Saito, Toshio Ohshima, Takaomi C. Saido, Hiroki Sasaguri

## Abstract

We previously developed single *App* knock-in mouse models of Alzheimer’s disease (AD) that harbor the Swedish and Beyreuther/Iberian mutations with or without the Arctic mutation (*App^NL- G-F^* and *App^NL-F^* mice). These models showed the development of amyloid β peptide (Aβ) pathology, neuroinflammation and cognitive impairment with aging. We have now generated *App* knock-in mice devoid of the Swedish mutations (*App^G-F^* mice) and some additional mutants to address the following two questions: [1] Do the Swedish mutations influence the mode of β-secretase inhibitor action *in vivo*? [2] Does the quantity of C-terminal fragment of amyloid precursor protein (APP) generated by β-secretase (CTF-β) affect endosomal properties as previously reported as well as other pathological events? Aβ pathology was exhibited by *App^G-F^* mice from 6 to 8 months of age, and was accompanied by microglial and astrocyte activation. We found that a β-secretase inhibitor, verubecestat, inhibited Aβ production in *App^G-F^* mice, but not in *App^NL-G-F^* mice, indicating that the *App^G-F^* mice are more suitable for preclinical studies of β-secretase inhibition given that most AD patients do not carry Swedish mutations. We also found that the quantity of CTF-β generated by various *App* knock-in mutants failed to correlate with endosomal alterations or enlargement, implying that CTF-β, endosomal abnormalities, or both are unlikely to play a major role in AD pathogenesis. This is the first AD mouse model ever described that recapitulates amyloid pathology in the brain without the presence of Swedish mutations and without relying on the overexpression paradigm. Thus, experimental comparisons between different *App* knock-in mouse lines will potentially provide new insights into our understanding of the etiology of AD.

## INTRODUCTION

Alzheimer’s disease (AD), the most prevalent cause of dementia, has been intensively investigated worldwide for over 100 years since it was first reported ^1^. There are currently, however, no efficacious disease-modifying treatments available for AD, although aducanumab^2^, an anti-Aβ human monoclonal antibody, was approved for use by the U.S. Food and Drug Administration in June 2021 following positive Phase 4 trial outcomes. To date, significant research advances have been achieved thanks to mouse models that recapitulate aspects of the AD pathophysiology seen in humans. Most AD mouse models overexpress mutant amyloid precursor protein (APP) or APP/presenilin 1 (PS1) cDNAs inserted into unknown loci of the host animals, which causes artificial aspects of their complex phenotypes ^3^. We previously developed *App^NL-F^* and *App^NL-G-F^* knock-in mice that harbor the Swedish (KM670/671NL) and Beyreuther/Iberian (I716F) mutations – with or without the Arctic (E693G) mutation – that do not depend on APP or APP/PS1 overexpression for their pathophysiological phenotype. These *App* knock-in mice exhibit age-dependent neuritic plaques composed of amyloid β peptide (Aβ) in the brain, followed by gliosis and memory impairment ^4^.

It should be noted, however, that the Swedish mutations, located adjacent to the cleavage site of APP by β-secretase, results in a drastic increase in CTF-β levels and influences the *in vitro* APP processing efficacy of β-secretase inhibitors^5^. The presence of Swedish mutations therefore renders the *App^NL-F^* and *App^NL-G-F^* lines as unsuitable for preclinical studies of β-secretase inhibitors. In effect, Swedish mutations are present in most APP transgenic mouse models that overexpress APP, and moreover, there is no single *App* knock-in mouse model that recapitulates amyloid pathology in the brain in the absence of Swedish mutations. In addition, recent cell-based studies have reported that CTF-β, not Aβ, contributes to early endosomal dysfunction^6 7^. Although multiple lines of evidence indicate that aberrant events in the endosomal trafficking system may appear as a common cytopathology regardless of whether the AD is early- or late-onset ^8, 9^ ^10^ ^11^ ^12^, it is not well understood if CTF-β affects early endosomal dysfunction *in vivo*.

In this study, we used a CRISPR/Cas9 system to develop *App^G-F/G-F^* knock-in (*App^G-F^*) mice harboring the Arctic and Beyreuther/Iberian mutations but devoid of the Swedish mutations ^13, 14^. Similar to the *App^NL-F^* and *App^NL-G-F^* lines, the *App^G-F^* line showed an age-dependent amyloid pathology, neuroinflammation and synaptic alteration. Acute administration of verubecestat ^15, 16^, a potent selective BACE1 inhibitor, reduced Aβ levels in *App^G-F^* mice, but not in *App^NL-G-F^* mice. We also found that early endosomal enlargement was present in the brains of *App^G-F^* mice even though the CTF-β quantity was quantitatively comparable to that of WT mice. Our findings demonstrate that BACE1 activity can be appropriately evaluated in *App^G-F^* mice without the interference of the Swedish mutations and that endosome enlargement does not correlate with CTF-β levels *in vivo*.

## RESULTS

### Generation of *App^G-F^* and *App^huAβ^*Mice By CRISPR/Cas9

We previously developed *App^NL-G-F^* mice by manipulation of the mouse *App* gene using a knock-in strategy^4^. Exon 16 of the *App* gene contains the Swedish mutations (KM670/671NL) while exon 17 contains the Arctic and Beyreuther/Iberian mutations (**Figure 1A**). Firstly, single-guide RNA (sgRNA)-App-Exon16 and single-stranded oligodeoxynucleotide (ssODN) containing the WT sequence to substitute the Swedish mutations (NL670/671KM) together with Staphylococcus aureus Cas9 (SaCas9) mRNA, where the proto-spacer adjacent motif (PAM) sequence is required as NNGRRT, were injected into the cytoplasm of heterozygous zygotes of *App^NL-G-F^* mice. The PAM sequence overlapped with the Swedish mutations so that, when knock-in of the WT sequence occurred, it could prevent sequential cleavages by SaCas9 because the original PAM site had disappeared (**Figures 1A and 1B**). Sanger sequencing analysis revealed that the desired substitution via homology-directed repair occurred successfully in the *App^NL-G-F^* allele of the founder mice with an efficiency of 10.8% (**Figure 1C**). Crossing the founder mice with WT mice to generate F1 mice, we confirmed that the Swedish mutations were fully removed from the *App^NL-G-F^* allele (**Figure 1C**). Using an identical strategy in *App^NL^* zygotes (see methods), we also generated *App^huAβ^*mice that carry only the humanized Aβ sequence in without any familial AD-causing mutation. We confirmed that there were no unexpected mutations in exons 16, 17 and 18 of the *App* gene in *App^G-F^*, *App^huAβ^*mice and others (**Figure S1**), indicating that all these lines are isogenic. *App^G-F/wt^* mice were then intercrossed to obtain homozygous *App^G-F/G-F^* mice that were viable. To explore the off-target effects of CRISPR/Cas9-mediated genome editing in the founder mice, we searched for potential off-target sites using the online tool COSMID ^17^ and Cas-OFFinder ^18^ (**Figure 1D**). Targeted sequencing analysis focusing on the candidate genomic regions revealed that no off-target modification took place in the founder mice of *App^G-F^* and *App^huAβ^*mice.

**Figure 1.**
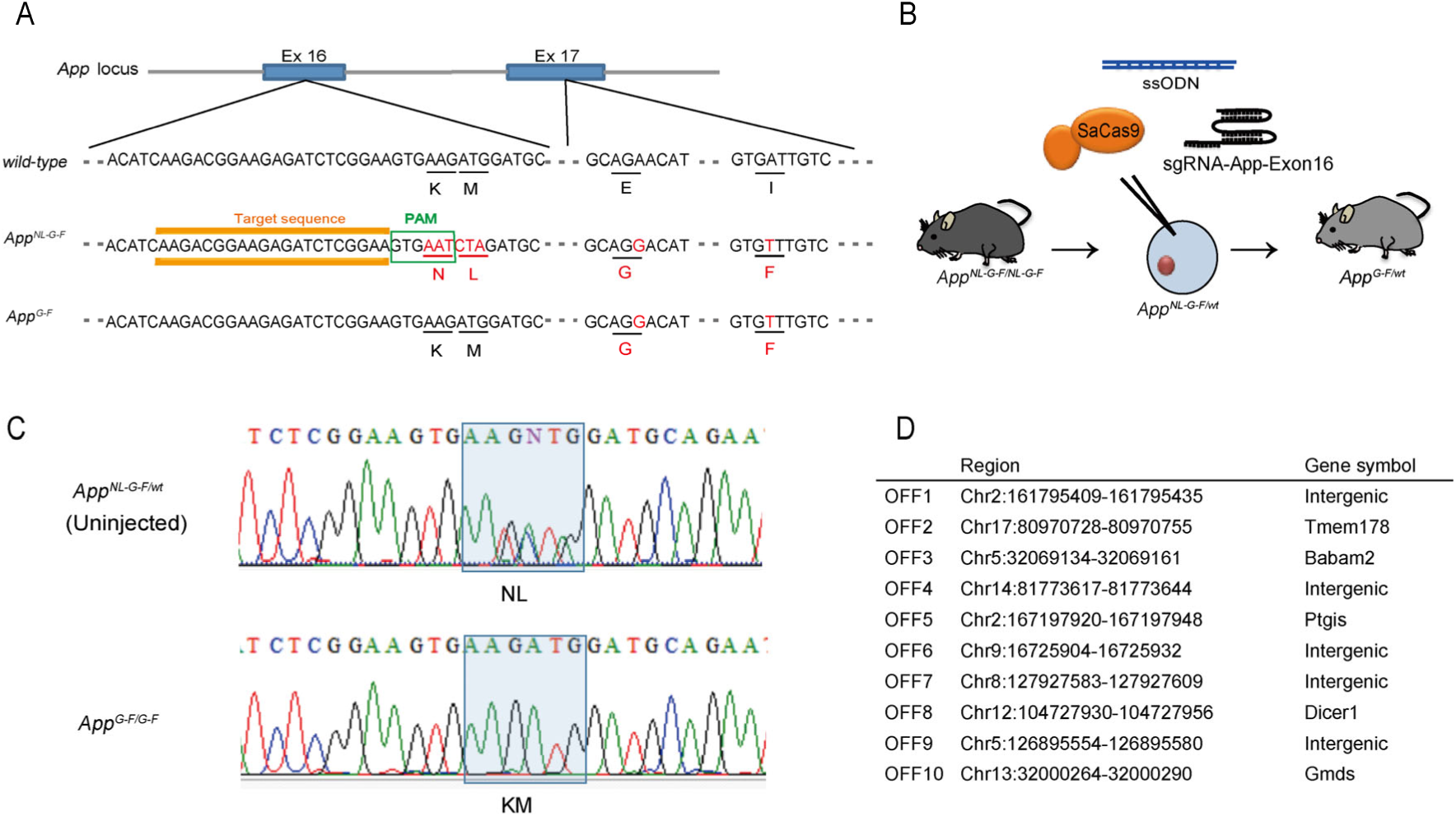
Generation of the single *App^G-F^* knock-in mice. (A) Exact sequences showing sgRNA (orange) with the PAM site (green) in the mouse *App* gene. Red characters represent the Swedish (KM670/671NL), Arctic (E693G) and Beyreuther/Iberian (I716F) mutations, respectively. (B) Schematic illustration of CRISPR/Cas9-mediated genome editing in *App^NL-G-F^* knock-in mouse zygotes by microinjection. (C) Sanger sequencing results determined *App^NL-G-F/wt^* (upper panel) and *App^G-F/G-F^* genotype (lower panel). The desired mutation loci (NL670/671KM) are indicated as a rectangular shape in blue shading. See also Figure S1. (D) Regional information of potential off-target sites which were identified using Cas-OFFinder (http://www.rgenome.net/cas-offinder/) and COSMID (https://crispr.bme.gatech.edu/).

### Neuropathology of *App^G-F^* Mice

We next analyzed the extent of amyloid pathology in the *App^G-F^* mice. Aβ_42_ levels in the cortex were age-dependently increased in the Tris-HCl and Guanidine-HCl (GuHCl) soluble fractions, with Aβ_40_ levels remaining relatively stable (**Figures 2A and 2B**). We also observed that progressive amyloid pathology mainly in the cortex and hippocampus occurred in an age-dependent manner (**Figures 2C and 2D**). Initial deposition of Aβ was observed around 4 months of age in the *App^G-F^* mice. At 12 months, Aβ deposition in the brains of *App^G-F^* mice detected in a much larger area than that in the *App^NL-F^* mice, but at a lower level to that in *App^NL-G-F^* mice (**Figure S2**). In addition, we analyzed the Aβ species constituting amyloid plaques in the *App^G-F^* mice using N-, C-terminal (Aβ_40,_ Aβ_42_) and Aβ_3(pE)-X_ (pE: pyroglutamate) specific antibodies. Aβ_40,_ Aβ_42_ and Aβ_3(pE)-X_ species were detected in the brain with a predominant deposition of Aβ_42_ over Aβ_40_ (**Figure 2E**). These results are consistent with the neuropathology observed in sporadic AD patients and in *App^NL-F^*and *App^NL-G-F^* mice ^4^.

**Figure 2.**
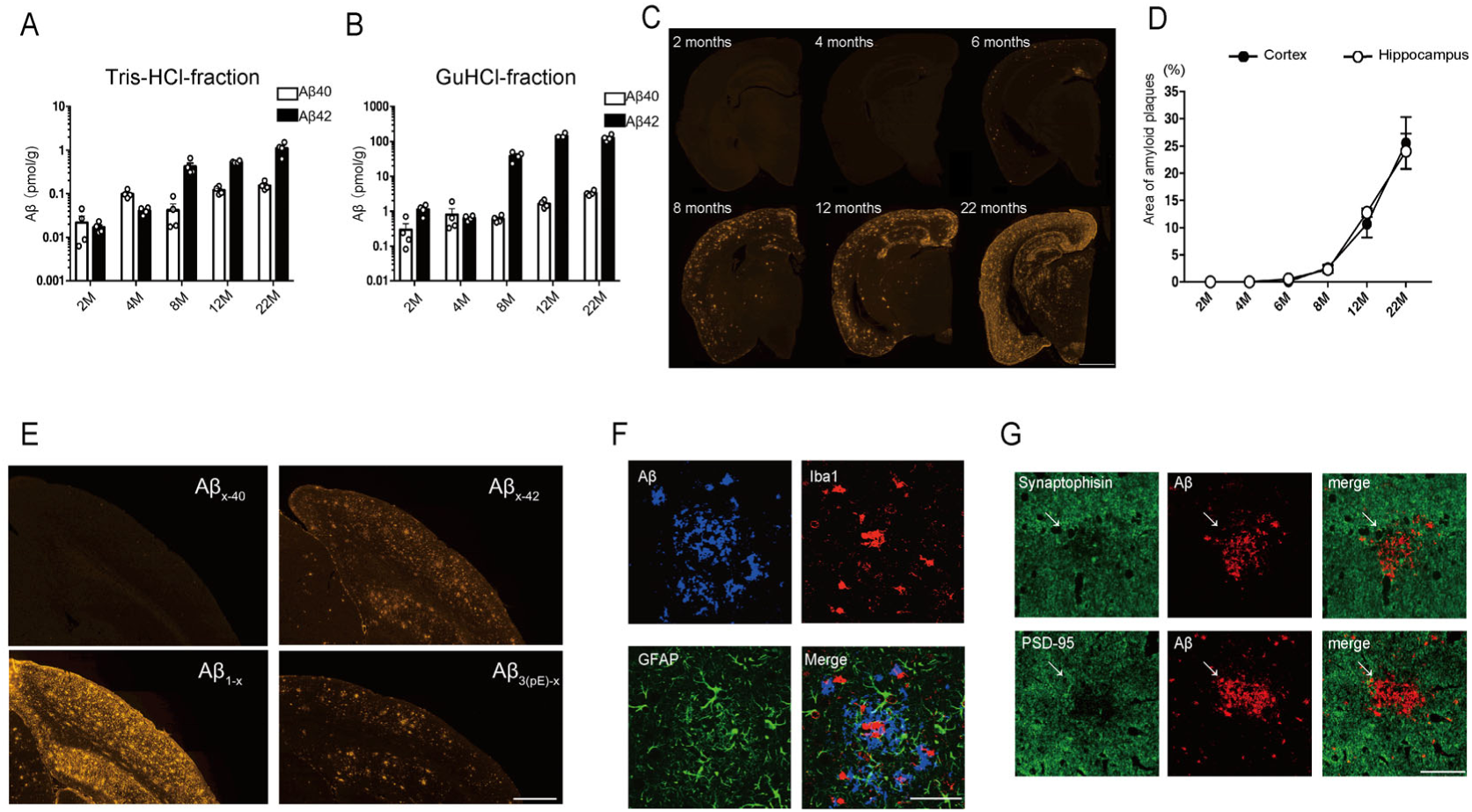
Neuropathology of *App^G-F^* mice. (A-B) Aβ content detected by ELISA using Tris-HCl soluble fraction (A) and GuHCl soluble fraction (B) of the cortices of *App^G-F^* mice at 2- 4-, 8-, 12- and 22 months (n=4 at each time point). Each bar represents the mean ± SEM. (C) Immunohistochemistry images showing Aβ deposition as indicated by immunostaining with N1D antibody against Aβ_1-5_. Scale bars indicate 1 mm. (D) Quantitative analysis of amyloid plaque areas in the cortices and hippocampi of *App^G- F^* mice at 2-, 4-, 8- and 12 months (n=4 at each time point) and at 22 months (n=3). Each bar represents the mean ± SEM. (E) Specific antibodies against N- (Aβ_1-x_ and Aβ_3(pE)-x_) and C- (Aβ_x-40_ and Aβ_x-42_) terminus of Aβ reveal the deposition of each species of Aβ in the brains of 22- month-old *App^G-F^* mice. Scale bars represent 500 µm. (F) Inflammatory responses in the cortices of *App^G-F^* mice at 22 months. Astrocytes (green) and microglia (red) can be seen surrounding Aβ (blue), as detected by triple staining with antibodies against GFAP, Iba1 and the N-terminus of human Aβ (82E1), respectively. Scale bars represent 100 µm. (G) Synaptic alteration detected in the hippocampus of a 22-month-old *App^G-F^* mouse. Aβ detected by 4G8 antibody against Aβ_17-24_was double stained with synaptophysin antibody as a presynaptic marker or with PSD95 antibody as a postsynaptic marker. White arrows indicate synaptic loss near Aβ aggregation. Scale bars represent 25 µm.

Chronic inflammation surrounding Aβ plaques in the brain is a pathological hallmark of AD. We therefore investigated the status of glial cells surrounding amyloid plaques in the *App^G- F^* mice. Reactive astrocyte and activated microglia are pathological signs of neuroinflammation, with evidence of both being observed (**Figure 2F**). We also examined pre- and post-synaptic alterations in brain slices and detected loss of synaptophysin and PSD-95 immunoreactivity near the Aβ plaques, which is consistent with those in other *App* knock-in mice (**Figure 2G**)^4^.

### Assessment of BACE1 Inhibition in *App^G-F^* Mice

The Swedish mutations have been considered to underlie the decreased APP processing potency of β-secretase inhibitors. Previous studies have shown that β-secretase inhibition is less efficacious in cells stably overexpressing the *APP*-containing Swedish mutations than from cells transfected with wild-type APP ^19, 20^. To compare the potency of β-secretase inhibition in animal models – with or without Swedish mutations in the *APP* gene – not relying on the overexpression paradigm, we administrated verubecestat, a potent BACE1 inhibitor, to 3-month-old WT, *App^NL- G-F^* and *App^G-F^* mice following a previously reported experimental protocol ^16^. We found that a single oral administration of verubecestat at the dose of 10mg/Kg significantly reduced both Aβ_40_ and Aβ_42_ levels in the cortices of *App^G-F^* mice, but not in *App^NL-G-F^* mice, 3 hours after treatment (**Figures 3A-3D**). These results indicate that the Swedish mutations are responsible for the poor potency of BACE1 inhibitors *in vivo* and that *App^G-F^* mice could serve as a powerful tool for the precise characterization of BACE1 and candidate inhibitory compounds.

**Figure 3.**
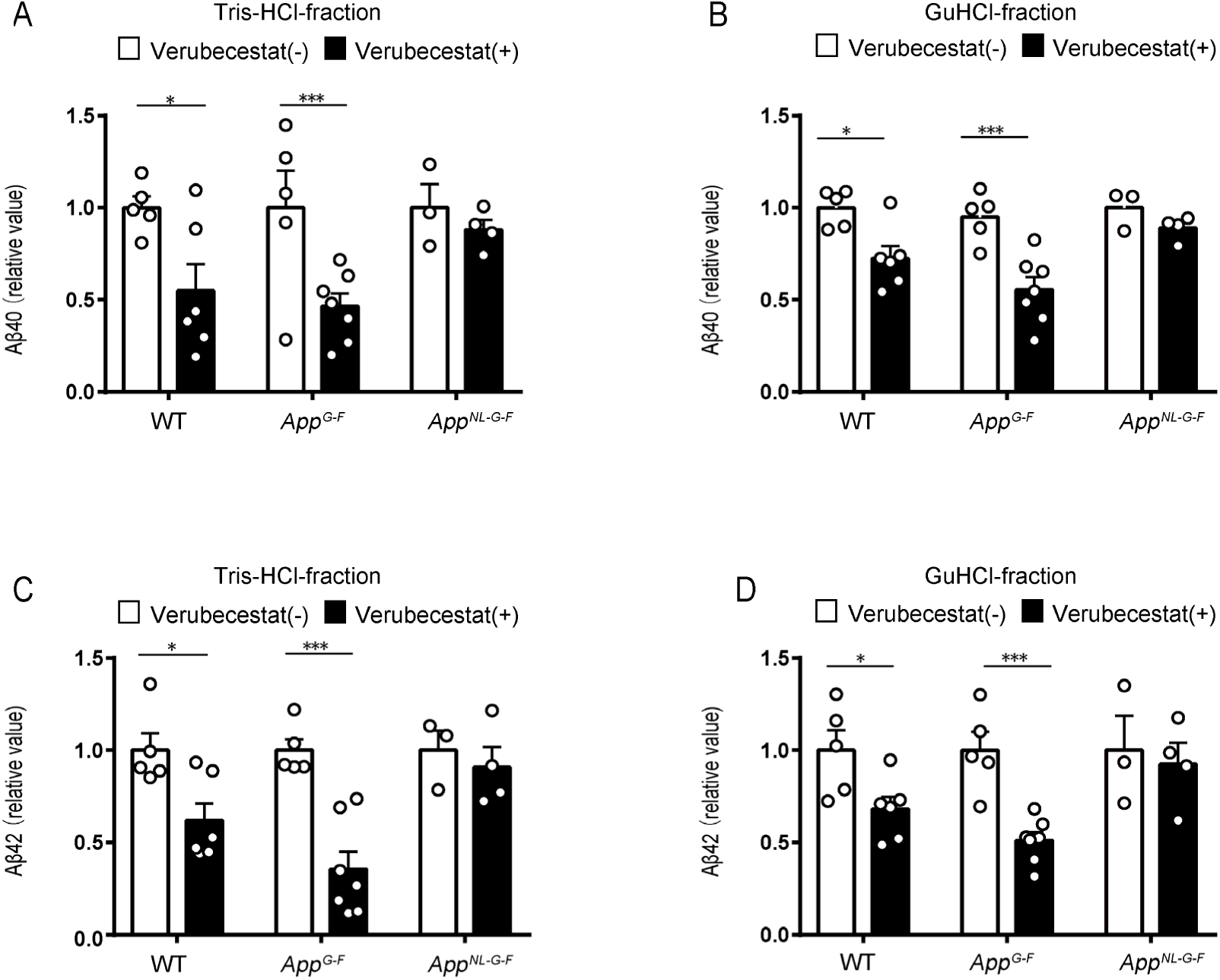
Removal of the Swedish mutations rescues the BACE1 inhibitory effect of verubecestat. (A-D) Aβ_40_ and Aβ_42_ levels detected by ELISA were decreased both in the Tris-HCl fraction (A, C) and GuHCl fraction (B, D) of 3-month-old wild-type and *App^G-F^* mice, but not in *App^NL-G-F^* mice. Each bar represents the mean ± SEM. * *P* < 0.05, ** *P* < 0.01, *** *P* < 0.001. (WT; verubecestat (+) n=5, (-) n=6, *App^G-F^*; (+) n=5, (-) n=7 and *App^NL-G-F^*; (+) n=3, (-) n=4, Student’s t-test).

### Relationship Between the Quantity of CTF-β and Endosomal Abnormality *in vivo*

Several studies have reported that the Swedish mutations alter the APP processing and shift the processing towards an amyloidogenic pathway via a competitive behavior between α- and β- secretases ^5, 21^. In an earlier study, we showed that the ratio of CTF-β/α levels in *App^NL-F^*and *App^NL- G-F^* mice is higher than that in wild-type mice ^4^. Here, we used five different *App* knock-in lines and WT mice to examine the effect of the mutations on the quantity of CTF-β and the extent of endosomal dysfunction (**Table 1**). The ratio of CTF- β/α in *App^G-F^* mice was much lower compared to *App^NL^*, *App^NL-F^* and *App^NL-G-F^* mice with no alteration of APP levels (**Figures 4A and S3A-S3E**). This finding indicates a slight shift to the β-cleavage pathway. We next examined whether CTF-β affects endosomal function *in vivo*. Some groups suggest that accumulated CTF-β itself induces endosome abnormalities independent of Aβ toxicity ^6, 7^. The quantity of CTF-β in the brains of *App^G-F^* mice was comparable to that of WT controls (**Figures 4A and 4B**). We focused on early endosomal antigen 1 (EEA1) as an early endosome marker and performed immunohistochemical analyses of the hippocampal CA1 region in six mouse lines: WT, *App^huAβ^*, *App^NL^*, *App^NL-F^*, *App^G-F^* and *App^NL-G-F^* knock-in mice. We detected a significant increase in the mean EEA1⁺ area in the CA1 pyramidal cell layer of five mutant lines compared with that of WT mice (**Figures 4C and 4D**). Kwart *et al*. demonstrated that endosome enlargement is a common pathology in familial APP mutant iPSC neurons, and that the quantity of CTF-β correlates with endosome abnormality. We consistently observed a significant alteration of the distribution of endosome size in the five mutant mouse lines, including *App^huAβ^*knock-in mice, compared with that of wild-type mice (**Figure 4E**). This was seen as an increase in the ratio of larger endosomes (>1 µm^2^) and a decrease in the ratio of smaller endosomes (<0.5 µm^2^) (**Figure 4E**). However, CTF-β levels did not correlate with endosome enlargement. Of note, endosomal sizes were enlarged in the brains of *App^G-F^* mice, the extent of which was similarly observed in *App^NL-G-F^* mice irrespective of large differences in CTF-β levels (**Figures 4B and 4F**). We sequenced the C-terminal region of APP in our *App* knock-in mouse lines and excluded the possibility that unexpected mutation was causing this alteration (**Figure S1**). These findings suggest that early endosomal enlargement might be caused by not only CTF-β but also by other toxic agents including human Aβ secretion in mouse brains.

**Figure 4.**
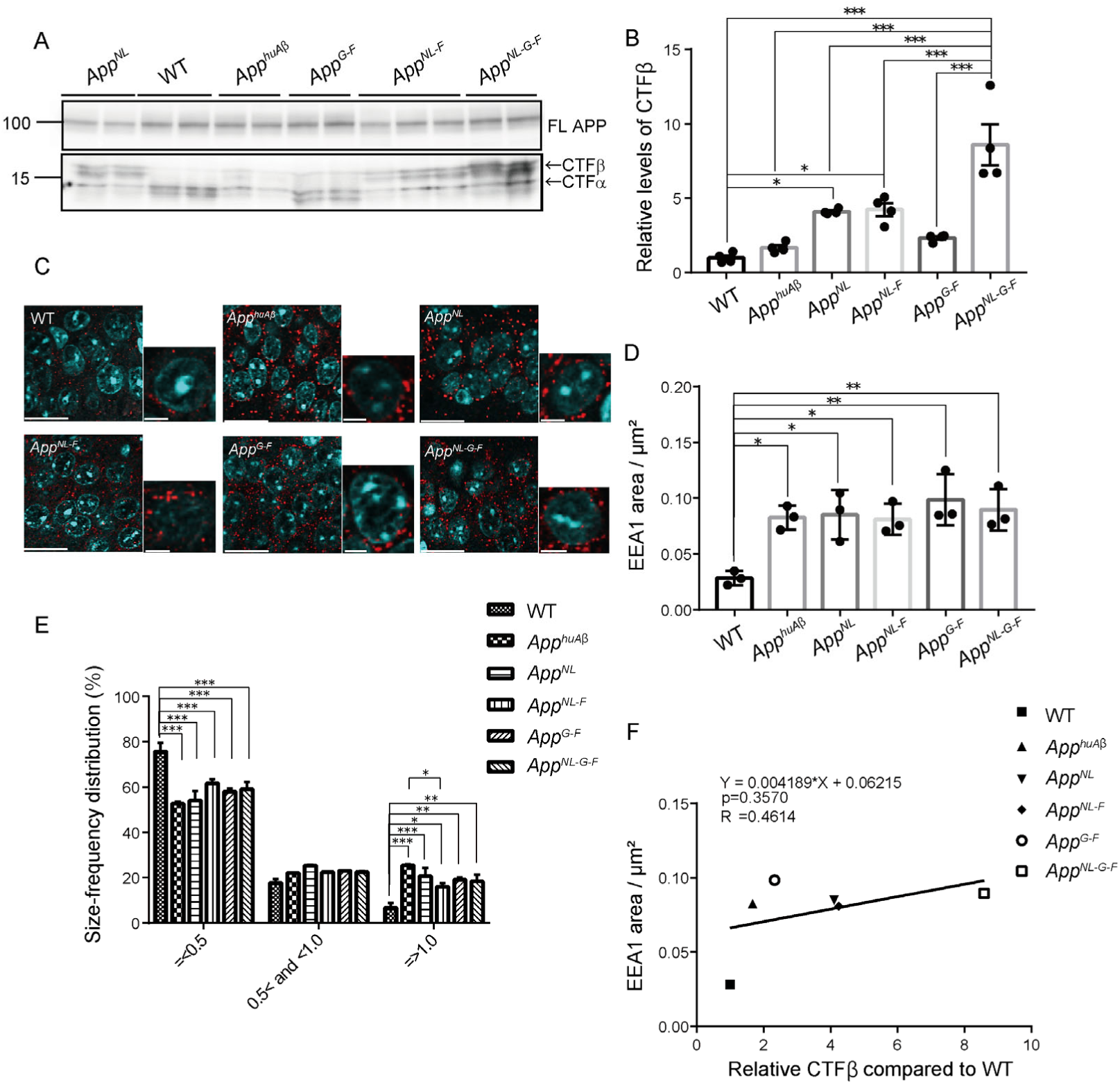
APP CTF expression levels and endosome abnormalities. (A) APP CTF expression in the brains of 12-month-old WT, *App^huAβ^*, *App^NL^*, *App^NL-F^*, *App^G-F^* and *App^NL-G-F^* mice. See also Figures S3-S4. (B) Quantification of relative levels of APP CTF-β relative levels compared to WT. WT, *App^huAβ^*, *App^NL^*, *App^NL-F^*, *App^G-F^* and *App^NL-G-F^*; n=4 for each genotype, one-way ANOVA followed by Tukey’s multiple comparison test. (C) Immunohistochemical images of early endosomes in CA1 pyramidal cells detected by EEA1 antibody (red) and Hoechst33342 staining of nuclei (blue). Brain sections from 12-month-old WT, *App^huAβ^*, *App^NL^*, *App^NL-F^*, *App^G-F^* and *App^NL-G-F^* mice. Scale bars indicate 20 µm (left) and 5 µm (right), respectively for each genotype. (D) Statistical analysis of EEA1^+^ area per µm^2^ in pyramidal cells of hippocampal CA1 region. (E) Endosomal size distribution was statistically analyzed using MetaMorph imaging software. WT, *App^huAβ^*, *App^NL^*, *App^NL-F^*, *App^G-F^* and *App^NL-G- F^*; n=3 for each genotype, one-way ANOVA followed by Tukey’s multiple comparison test (D and E). Each bar represents the mean ± SEM. * *P* < 0.05, ** *P* < 0.01, *** *P* < 0.001 (B, D and E). (F) Relationship between EEA1 area / µm^2^ and CTF-β content. Pearson’s correlation coefficient R, the associated p value and linear regression equation are shown in the figure.

**Table 1.**
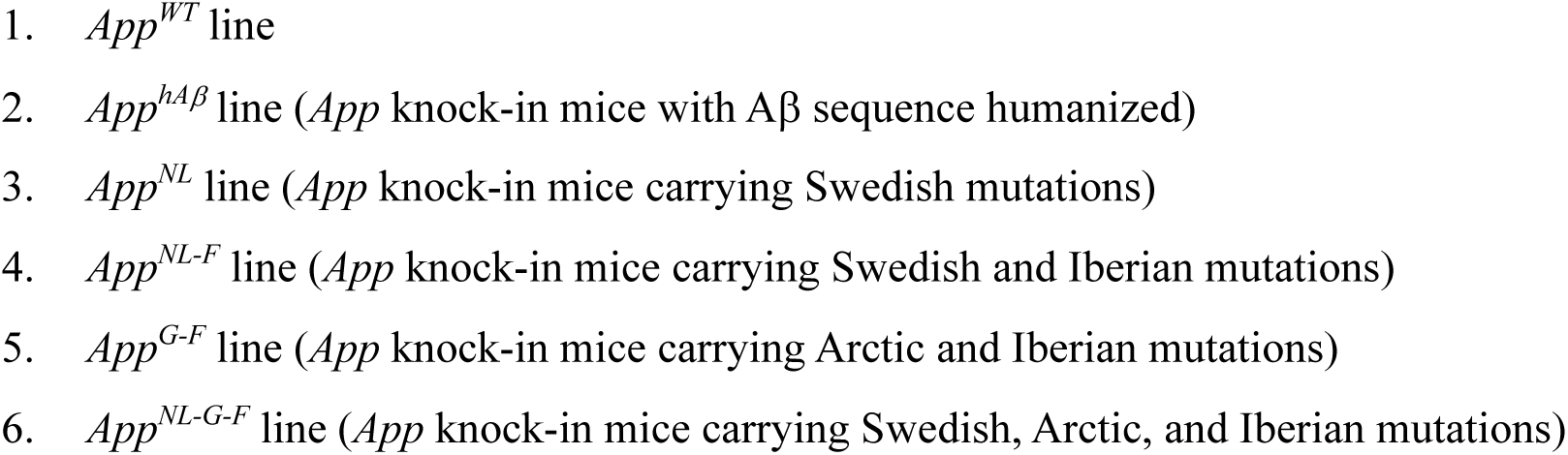
Isogenic panel of mouse lines used in the present study

## DISCUSSION

In the present study, by removing the Swedish mutation from *App^NL-G-F^* mice we developed a new *App* knock-in line, *App^G-F^*, which harbors both the Arctic and Beyreuther/Iberian mutations. The *App^G-F^* mice exhibited an age-dependent and typical amyloid pathology, neuroinflammation, characterized by reactive astrocytes and activated microglia surrounding the Aβ plaques, and aberrant pre- and post-synaptic structures near the plaques. Verubecestat intervention effectively reduced Aβ levels in the cortices of *App^G-F^* mice, but not in the conventional *App* knock-in mice containing the Swedish mutation. Endosomal enlargement was also observed, although the CTF-β levels in the brains of *App^G-F^* mice was comparable to those of WT mice.

Aβ deposition in the brains of *App^G-F^* mice occurs from around 4 months of age, compared to around 2 months of age in *App^NL-G-F^* mice and 6 months in *App^NL-F^* mice ^4^, suggesting that the *App^G-F^* mice serve as “a moderate model” of the three lines from the point of view of amyloidosis in the mouse brain (**Figure S2**). *App^G-F^* mice also showed an age-dependent amyloid pathology in the subcortical area as well as in the cortex and hippocampus, which is consistent with human carriers of the Arctic mutation ^22^. This is the first AD mouse model that recapitulates amyloid pathology in the brain, but does not harbor the Swedish mutation and is not dependent on APP overexpression.

Previous studies based on transgenic mice overexpressing the *APP* gene with familial mutations and CRISPR/Cas9-mediated genomic modified iPSCs indicated that AD-associated early endosomal enlargement depends on the excess accumulation of CTF-β, but not Aβ ^6, 7, 23–27^. On the other hand, other studies have reported that Aβ toxicity is indeed a causative factor for impaired endocytic sorting ^28–30^. In this study, using *App* knock-in mice, we observed early endosomal abnormalities in hippocampal CA1 pyramidal neurons both with or without CTF-β overproduction compared to WT mice (**Figures 4 and S4**). Given that Aβ_42_ levels reached a plateau and that Aβ plaque formation was spread abundantly in the hippocampi of *App^G-F^* mice as young as 12 months of age (**Figures 2A and 2B**), Aβ species themselves, independent of CTF-β accumulation, are also likely to cause early endosomal enlargement *in vivo*. Future analyses on concerning which Aβ peptide(s) such as Aβ_40_, Aβ_42_, Aβ_3(pE)-X_ are responsible for altering the endocytic system are required to elucidate the underlying mechanism(s). Our results show that humanization of Aβ alone induces endosomal enlargement in mice, which is consistent with a transcriptomic study using weighed gene co-expression network analysis (WGCNA) indicating that Eea1 is one of the highly correlated genes explaining biological alterations in the hippocampi of huAβ-KI mice ^31^. On the other hand, recent studies indicate that endosomal enlargement occurs via an APP-independent pathway ^32 33^. Taken together, endosomal trafficking defects associated with AD cytopathogenesis may be occur via different means such as CTF-β-dependent, Aβ- dependent and APP-independent pathways.

A large number of BACE1 inhibitors have been explored and investigated as potent disease-modifying drugs in the AD research field, but all of them, to our knowledge, have failed to show efficacy in clinical trials. However, as the A673T (Icelandic) mutation^34^ positively established the proof-of-concept that the inhibition of β-secretase cleavage reduces the risk of AD onset, the discovery of BACE1 inhibitory compounds that pass through the blood-brain barrier and directly abrogate Aβ production in human brains remains a promising path to treat AD patients. Although single *App^NL-F^* and *App^NL-G-F^* knock-in mice have been used in more than 500 laboratories and pharmaceutical companies worldwide as second-generation mouse models of AD ^4^, these mice are not compatible with BACE1-related studies due to the presence of Swedish mutations. Our results provide consistent evidence that the Swedish mutations hinder the BACE1 inhibitory activities of verubecestat *in vivo*, similar to several reports showing the reduced activity of BACE1 inhibitors including not just verubecestat ^19^ but also other drug candidates in mice harboring the Swedish mutations ^35 21^. Thus, *App^G-F^* mice now profile as a novel type of single *App*-KI mice without the interference of the Swedish mutations. The potential exists for these mice to be used to efficiently and precisely to identify active compounds for BACE1 inhibition *in vivo* that might have been overlooked in a vast number of studies in which AD model mice were used that contained the Swedish mutations and were based on an APP overexpression paradigm. Our range of single *App* knock-in mice including the *App^NL-F^*, *App^NL-G-F^* and *App^G-F^* lines, are available for use by research groups and companies worldwide who can choose the AD mouse model line most suited to the purpose of their study.

## EXPERIMENTAL MODEL DETAIL

### Mice

All animal experiments were conducted in compliance with regulations stipulated by the RIKEN Center for Brain Science. *App^NL-F^* or *App^NL-G-F^* mice expressing two or three familial AD mutations [Swedish (KM670/671NL) and Beyreuther/Iberian (I716F) with or without the Arctic (E693G) mutation] driven by the endogenous promoter, as well as the humanized Aβ sequence, were generated described previously ^4^. *App^NL-G-F^* and ICR mice were used as zygote donors and foster mothers. C57BL/6J and *App^NL^* mice were prepared as controls^4^. All mutant mice used in this study were homozygous for the expressed mutations. Both male and female mice were used in our experiments. All mice were bred and maintained in accordance with regulations for animal experiments promulgated by the RIKEN Center for Brain Science.

## METHODS DETAILS

### Generation of *App^G-F^* mice

sgRNA targeting mouse *App* exon 16 was designed *in silico* utilizing the CRISPR design tool ^36^. To reduce the possibility of off-target events, SaCas9 that recognizes NNGRRT as the PAM site was selected to introduce double-stranded breaks. ssODN was designed to cause NL670/671KM substitution (AATCTA>AAGATG) overlapping the PAM region so that the oligonucleotide did not include silent mutations, thus preventing re-binding and re-cutting after the desired genome modification via homology -directed repair. A plasmid vector (Addgene, #61591) was used for *in vitro* transcription of SaCas9 mRNA, and sgRNA was synthesized as described previously ^37^. Information on the primers and oligonucleotides used for the *in vitro* synthesis of CRISPR tools is listed in **Table S1**. The prepared SaCas9 mRNA (100 ng/µl) and sgRNA (100 ng/µl) along with ssODN (100 ng/µl) were co-injected into the cytoplasm of *App^NL-G-F/wt^* zygotes. Founder mice were identified by PCR and sequencing analysis of the targeted site and crossed with wild-type mice to obtain heterozygous F1 mice.

### Generation of *App^huAβ^*mice

To generate *App^huAβ^*mice that carried only the humanized Aβ sequence, virtually the same strategy was used to that employed for developing *App^G-F^* mice. The prepared sgRNA, mRNA and ssODN were identical to those used for *App^G-F^* mice, with the only different being that *App^NL^* zygotes instead of *App^NL-G-F^* zygotes were used for injecting genome editing tools. Potential off-target sites were also identical as those for *App^G-F^* mice.

### Off-target effects analysis

Candidate sequences were identified *in silico* using COSMID (https://crispr.bme.gatech.edu/) ^17^ and Cas-OFFinder (http://www.rgenome.net/cas-offinder/) ^18^ allowing up to 3 bp mismatches and 1 bp DNA and/or RNA bulge. Genomic DNA extracted from mouse tails was amplified by PCR with the primers listed in **Table S2**. All genomic sequences of the amplicons were analyzed by Sanger sequencing using a DNA sequencer (ABI 3730xl).

### Genotyping

Genomic DNA was extracted from mouse tails in lysis buffer (10 mM pH 8.5 Tris-HCl, 5 mM pH 8.0 EDTA, 0.2% SDS, 200 mM NaCl, 20 µg/ml proteinase K) through a process of ethanol precipitation. Purified DNA was subjected to PCR and followed by Sanger sequencing analysis with the specific primers according to a previous report employing *App*-KI mice, including *App^NL^*, *App^NL-F^*and *App^NL-G-F^* strains ^4^. Genotyping primers for *App^G-F^* mice are listed in **Table S2**.

### Western blotting

Mouse brain tissues were homogenized in lysis buffer containing 50 mM Tris pH 7.6, 0.15 M NaCl, 1% Triton and cOmplete protease inhibitor cocktail (Roche Diagnostics) using a Multi-beads shocker (YasuiKikai). Homogenates were incubated at 4 °C for 1 h and centrifuged at 15000 rpm for 30 minutes, and the supernatants were collected as loading samples. Concentrations of protein samples were measured with the aid of a BCA protein assay kit (Thermo Fisher Scientific). Equal amounts of proteins were subjected to sodium dodecyl sulfate-polyacrylamide gel electrophoresis (SDS-PAGE) and transferred to PVDF membranes. For detection of APP-CTFs, delipidated samples were loaded onto membranes and boiled for 5 min in PBS before blocking with ECL primer blocking buffer (GE Healthcare). Membranes were incubated at 4 °C with primary antibodies against APP (MAB348, Millipore, 1:1000) or APP-CTFs (A8717, Sigma-Aldrich, 1:1000) with GAPDH (HRP-60004, Proteintech, 1:150000) as a loading control. Targeted proteins were visualized with ECL select (GE Healthcare) and a LAS-3000 Mini Lumino image analyzer (Fujifilm).

### Immunohistochemistry

Paraffin-embedded mouse brains were sectioned (thickness 4 µm) and subjected to deparaffinization processing; antigen retrieval was then performed by autoclaving at 121 °C for 5 min. Brain sections were treated with 0.3% H_2_O_2_ in methanol solution for 30 min to inactivate endogenous peoxidases. Sections were rinsed with TNT buffer (0.1 M Tris pH 7.5, 0.15 M NaCl, 0.05% Tween20), blocked using a TSA Biotin System kit and incubated at 4 °C overnight with primary antibodies diluted in TNB buffer (0.1 M Tris pH 7.5, 0.15 M NaCl). Primary antibody dilution ratios are listed in **Table S3**. Sections were washed, incubated with biotinylated secondary antibody and a tyramide signal amplification system used to detect amyloid pathology. For detection of neuroinflammatory signs and early endosome pathology, secondary antibodies conjugated with Alexa Fluor 555 diluted in TNB buffer or 0.2% casein in PBS were used. Sections were stained for 15 min with Hoechst33342 (Thermo Fisher Scientific) diluted in PBS, and then mounted with PermaFluor (Thermo Fisher Scientific). Section images were obtained using a confocal laser scanning microscope FV-1000 (Olympus) and a NanoZoomer Digital Pathology C9600 (Hamamatsu Photonics). Quantification of immunoreactive signals was performed using Metamorph Imaging Software (Molecular Devices) and Definiens Tissue Studio (Definiens).

### Enzyme-linked immunosorbent assay (ELISA)

Mouse brain samples were homogenized in lysis buffer (50 mM Tris-HCl, pH 7.6, 150 mM NaCl and protease inhibitor cocktail) using a Multi-beads shocker (YasuiKikai). The homogenates were centrifuged at 70000 rpm at 4°C for 20 min, and the supernatant was collected as a Tris-Soluble (TS) fraction to which 1/11 vol of 6 M guanidine-HCl (GuHCl) in 50 mM Tris and protease inhibitors were added. The pellet was loosened in lysis buffer with a Pellet Pestle (KIMBLE), dissolved in 6 M GuHCl buffer, and sonicated at 25 °C for 1 min. The sample was incubated for 1 hour at room temperature and then subjected to centrifugation at 70000 rpm at 25 °C for 20 min. The supernatant was collected as a GuHCl-soluble fraction. Tris-soluble and GuHCl-soluble fractions were applied to 96-well plates using an Aβ ELISA kit (Wako) according to the manufacturer’s instructions. For detection of Arctic Aβ produced from the brains of *App^NL-G-F^* and *App^G-F^* mice, standard curves were drawn using human Aβ peptides carrying the Arctic mutation as described previously ^4^.

### Verubecestat administration

Verubecestat (ChemScene) dissolved in PBS was administrated orally to 3-month-old mice using a flexible sonde (FUCHIGAMI) at a single dose of 10mg/kg according to Kennedy et al^16^. Three hours after a single treatment mouse brains were dissected and stored at -80 °C.

## QUANTIFICATION AND STATISTICAL ANALYSIS

All data are shown as the mean ±SEM within each figure. For comparisons between two groups, data were analyzed by Student’s- or Welch’s- *t*-test or Mann-Whitney test. For comparisons among more than three groups, we used one-way analysis of variance (ANOVA) followed by Dunnett’s post hoc analysis or Tukey’s post hoc analysis. All statistical analysis were performed using GraphPad Prizm 7 software (GraphPad software). Levels of statistical significance were presented as a *P*-values: * *P* < 0.05, ** *P* < 0.01, *** *P* < 0.001.

## AUTHOR CONTRIBUTIONS

NW, KS, KN, TS, TCS and HS designed the research plan. NW, KS, GS, AI, NK, MT, MS, NY, RF, NW, KI and ST performed the experiments. NW, KS, KN, TS, TCS and HS analyzed and interpreted data. HS, KS, NW, SH, TS and TCS wrote the manuscript together. HS, TO, TS and TCS supervised the entire research.

## ACKNOWLEDGEMENTS

We thank Nobuhisa Iwata, Nagasaki University, for valuable discussions. We also thank Yukiko Nagai-Watanabe for secretarial work. This work was supported by AMED under Grant Number JP20dm0207001 (Brain Mapping by Integrated Neurotechnologies for Disease Studies (Brain/MINDS)) (TCS) and JSPS KAKENHI Grant Number JP18K07402 (HS).

## SUPPLEMENTARY INFORMATION

**Figure S1.**
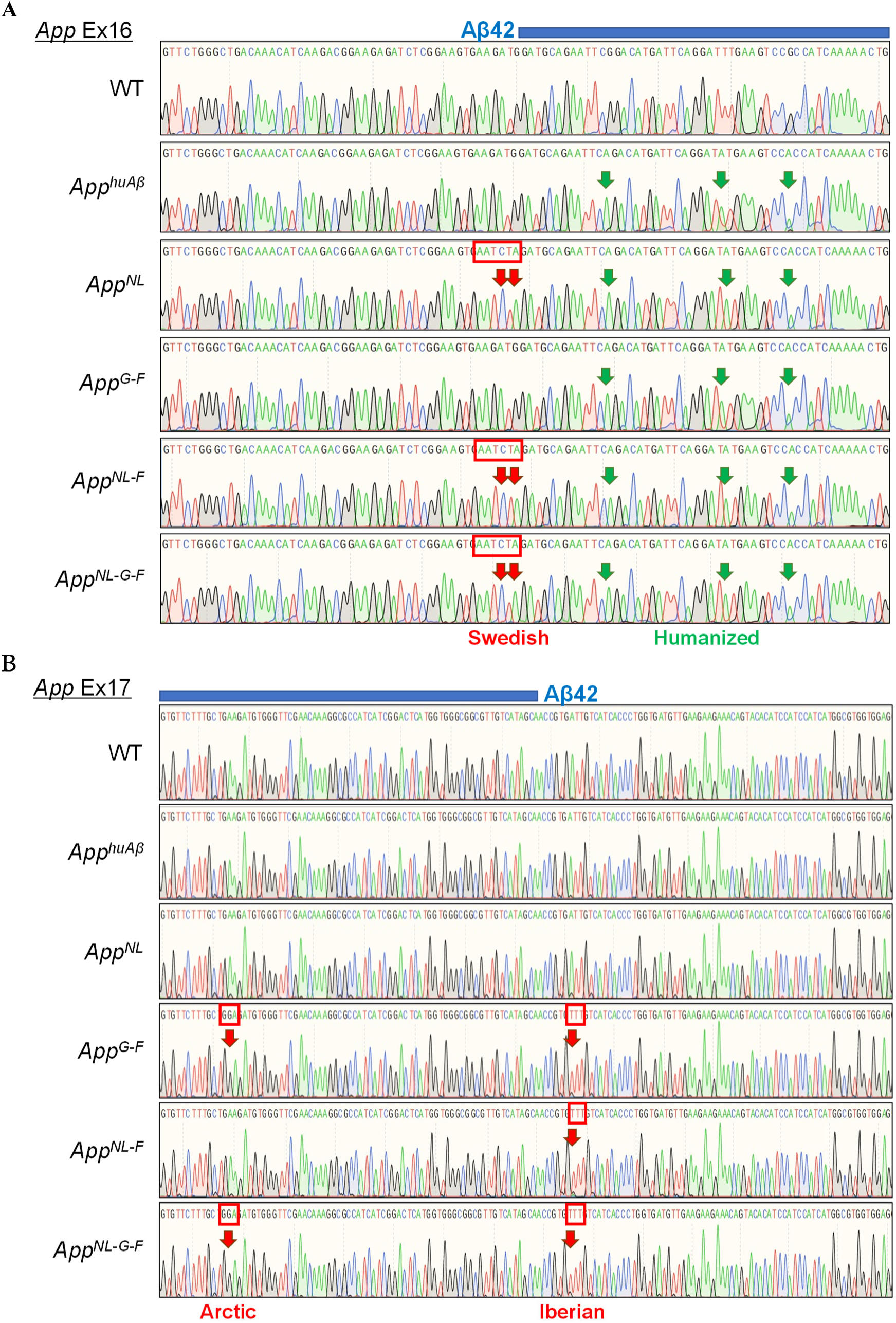
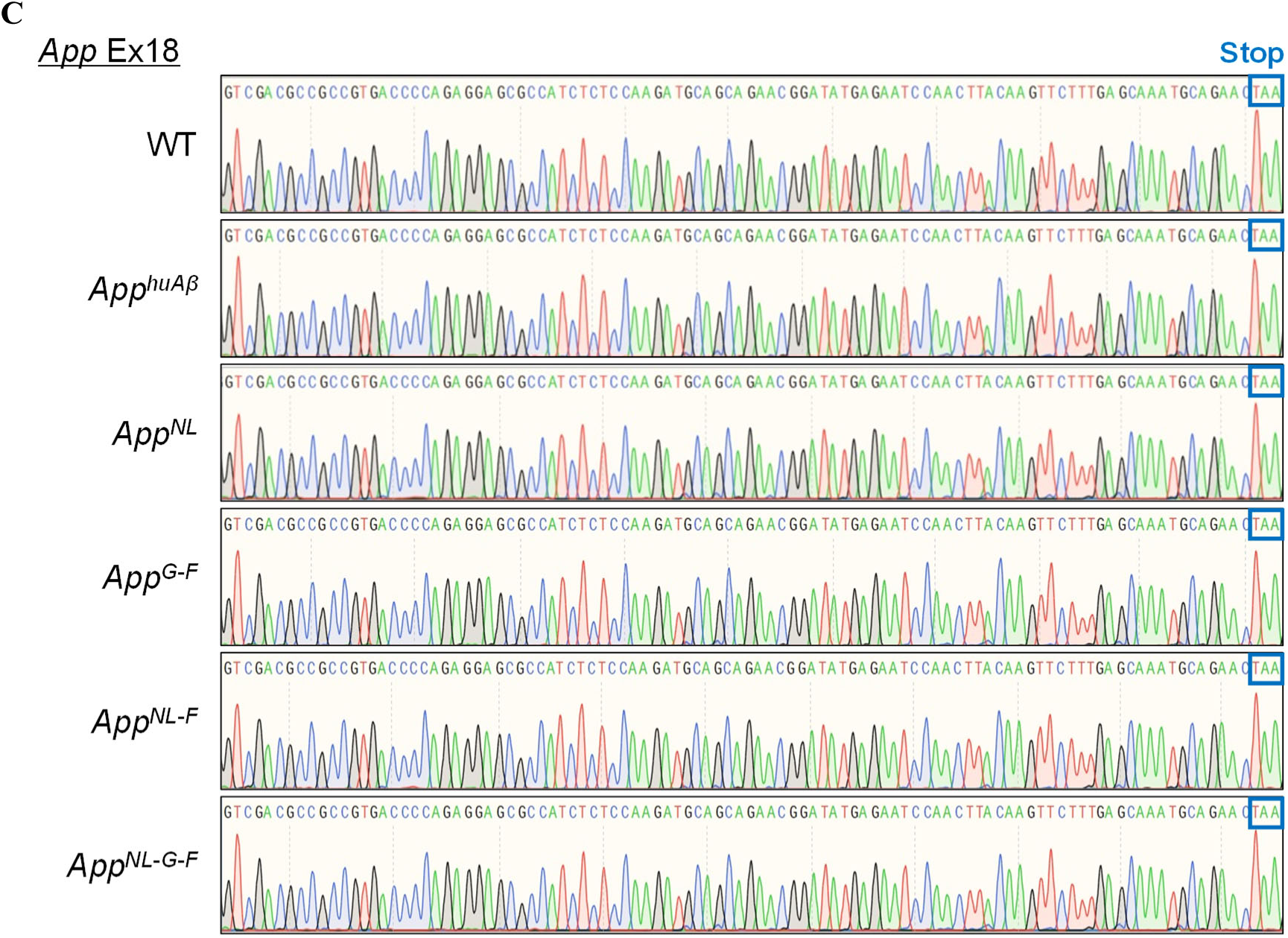
Exon sequences of the *App* gene in *App* knock-in mouse lines encoding CTF-β. Sequencing analyses of *App* exon 16 (A), exon 17 (B) and exon 18 (C) indicate that all genotypes of the *App* knock-in mice (Table 1) share identical amino acid sequences except for the artificially introduced mutations, indicating that the difference in the electrophoretic mobility of CTF-β between each line shown in **Fig. 4a** can be solely attributed to the intentionally altered mutations introduced.

**Figure S2.**
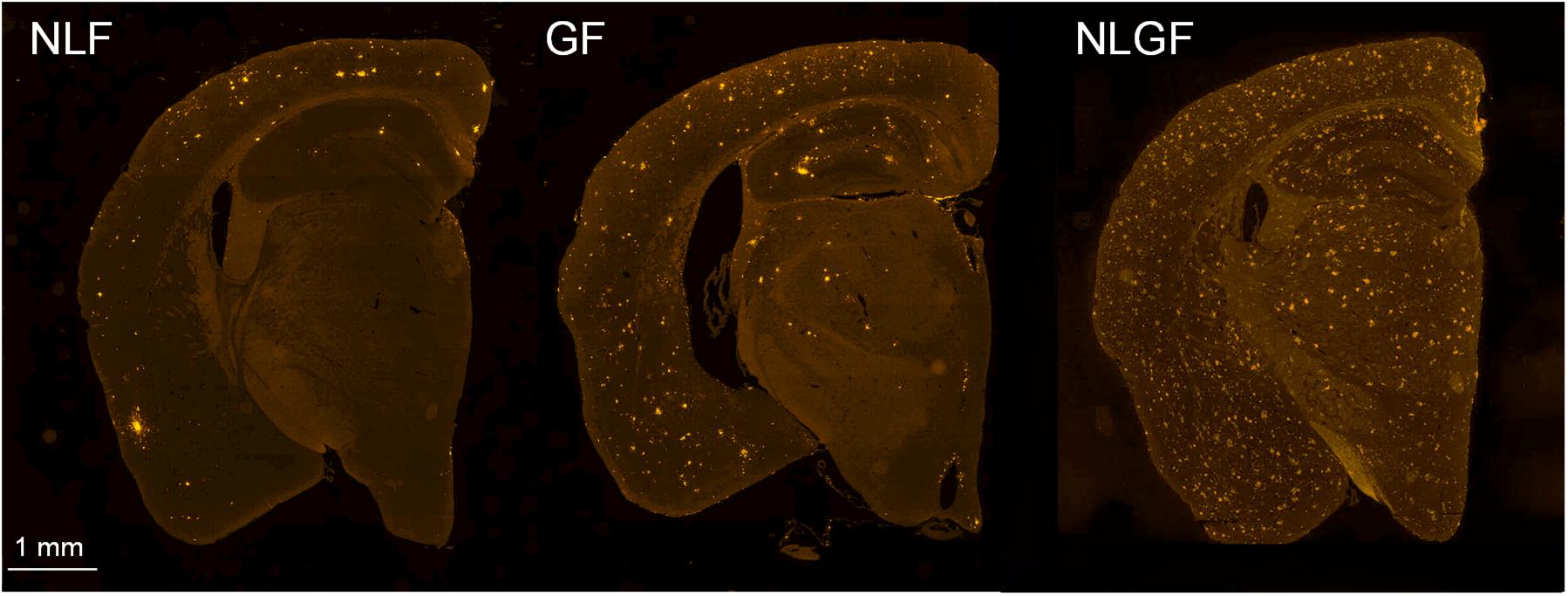
Aβ plaque formation in the brains of 12-month-old *App^NL-F^*, *App^G-F^* and *App^NL-G-F^* mice. Immunohistochemistry images with N1D antibody detection show that *App^G-F^* mice exhibited more and less prominent amyloid pathology than *App^NL-F^* and *App^NL-G-F^* mice, respectively, at 12 months.

**Figure S3.**
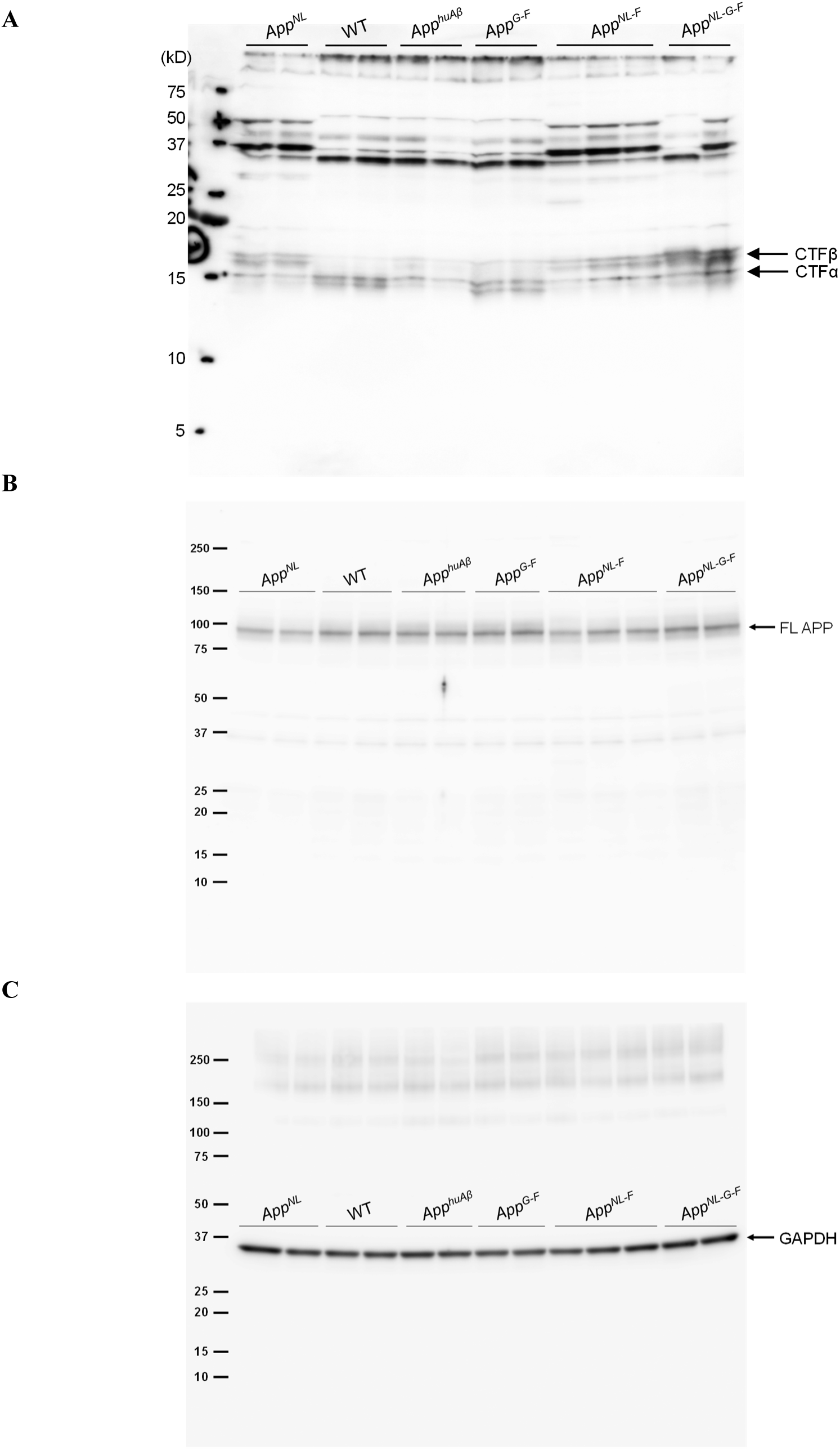
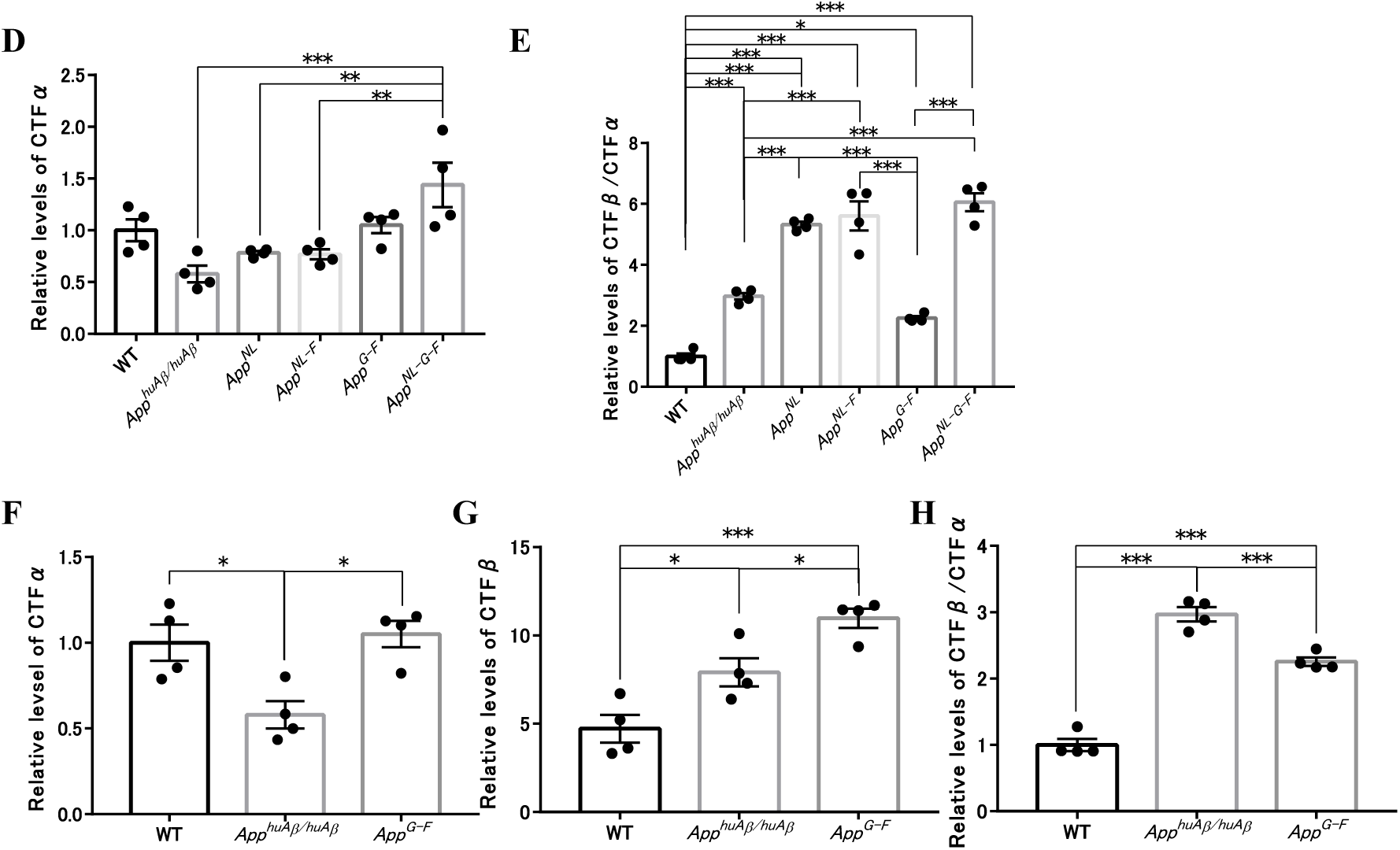
Western blot analysis of APP CTFs. (A-C) Full-blots of APP CTFs (A) and full-length APP (B) and GAPDH (C) using the cortices of 12- month-old *App*-knock-in lines used in **Figure 4** are shown. (D) Intensities of CTF-α-immunoreactive bands were quantified and statistically analyzed for WT, *App^huAβ^*, *App^NL^*, *App^NL-F^*, *App^G-F^* and *App^NL- G-F^* mice. (E) Relative levels of CTF-β/CTF-α ratio compared to WT mice. (E-G) Relative levels of CTF-α (F), CTF-β (G) and CTF-β/CTF-α ratios (H) were statistically analysed for WT, *App^huAβ^*and *App^G-F^* mice. Each bar represents the mean ± SEM. * *P* < 0.05, ** *P* < 0.01, *** *P* < 0.001. (WT; n=4, *App^huAβ^*; n=4, *App^NL^*; n=4, *App^NL-F^*; n=4, *App^G-F^*; n=4 and *App^NL-G-F^*; n=4, one-way ANOVA followed by Turkey’s multiple comparison test).

**Figure S4.**
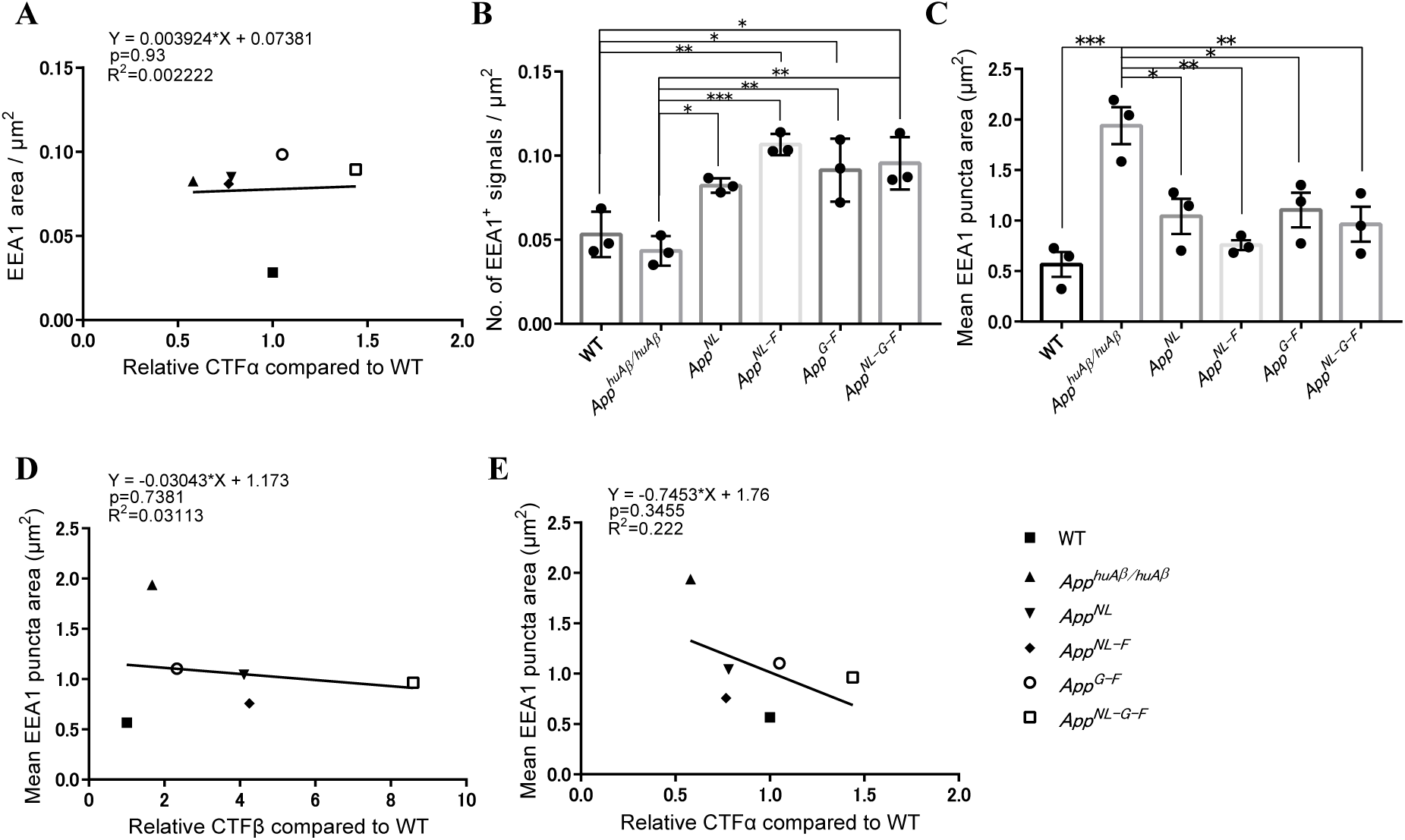
Analysis of APP CTFs and endosomal abnormalities. (A) CTF-α levels did not correlate with the area (µm^2^) of EEA1^⁺^. (B) Number of EEA1^+^ signals per µm^2^ was increased in App-knock-in mice except for *App^huAβ^*mice. (C) Mean EEA1 puncta area. (D-E) No correlation was observed between mean EEA1^+^ puncta area and levels of CTF-β (D) or CTF-α (E). Pearson correlation coefficient R and p value are shown along with the linear regression equation (A), (D) and (E). Each bar represents the mean ± SEM. * *P* < 0.05, ** *P* < 0.01, *** *P* < 0.001. (WT; n=3, *App^huAβ^*; n=3, *App^NL^*; n=3, *App^NL-F^*; n=3, *App^G-F^*; n=3 and *App^NL-G-F^*; n=3, one-way ANOVA followed by Turkey’s multiple comparison test) (B) and (C).

**Table S1.**
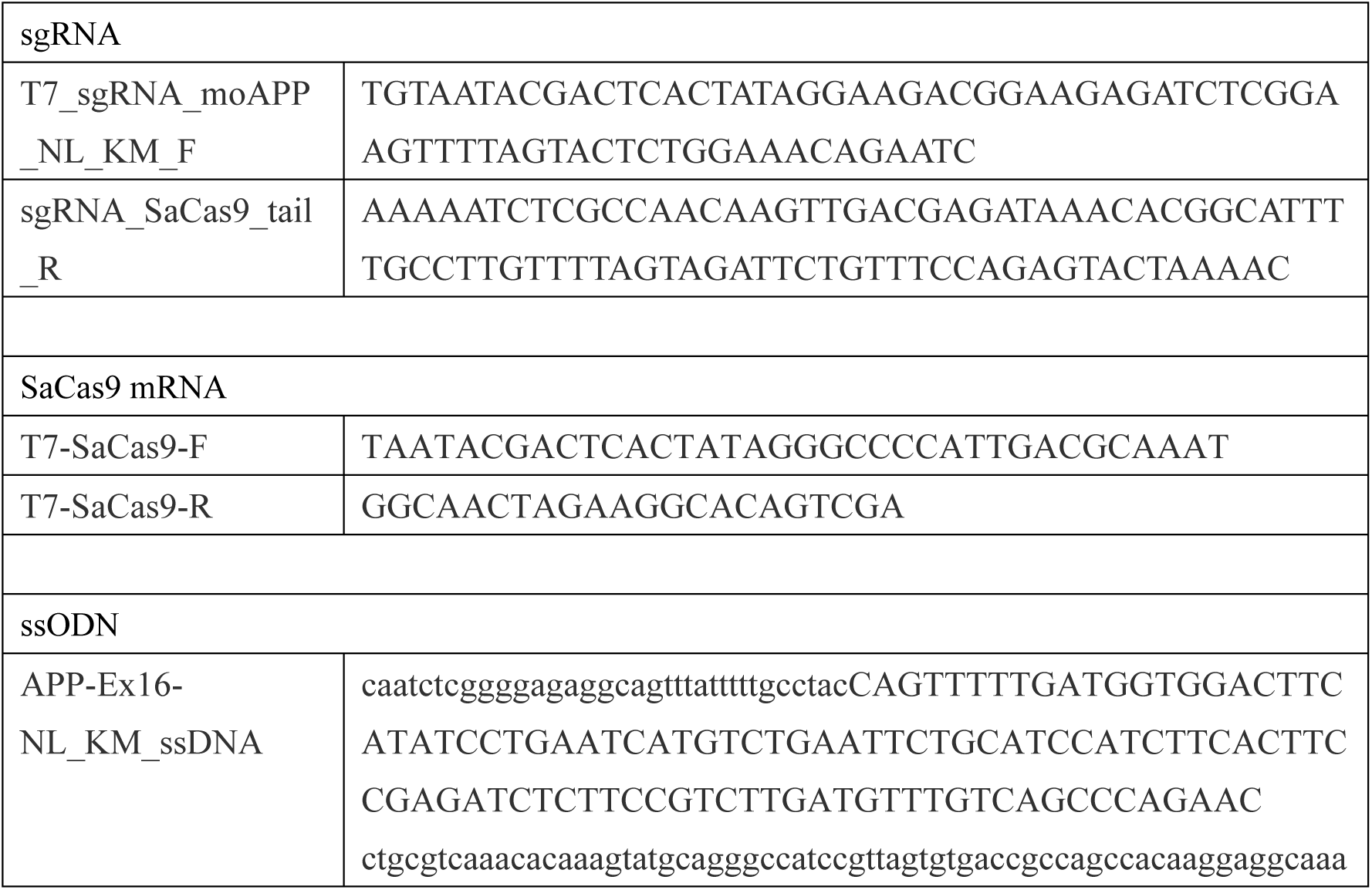
**Information on primers and oligonucleotides used for the synthesis of sgRNA, SaCas9 mRNA and ssODN.** In vitro synthesis of CRISPR tools was performed with the listed primers for the generation of *App^G- F^* and *App^huAβ^*mice. See Methods for details.

**Table S2.**
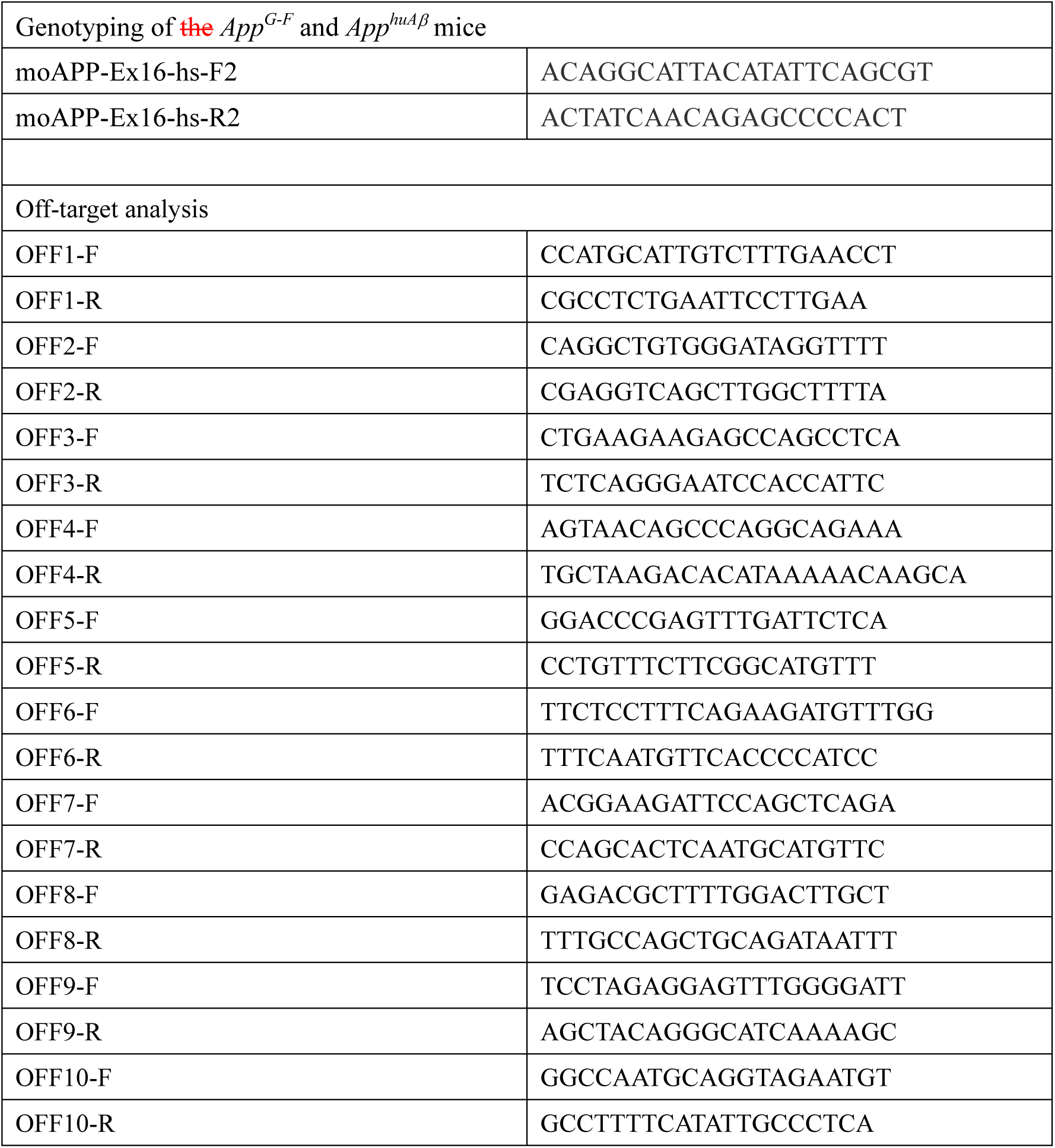
**List of primers used for genotyping and off-target analyses.** All primer pairs were used for PCR and subsequence sequencing analyses. The sequential number of OFF primers corresponds to that of the potential off-target sites shown in Figure 1. See Methods for details.

**Table S3.** **List of primary antibodies used for immunohistochemistry.** Primary antibody dilution ratios are shown. See Methods for details.

| Antibody | Dilution ratio | # Manufacturer or reference |
| --- | --- | --- |
| Aβ40 | 1:100 | #10047, IBL |
| Aβ42 | 1:100 | #18582, IBL |
| Aβ3pE | 1:400 | Saido et al., 1996 |
| N1D | 1:200 | Saido et al., 1996 |
| 82E1 | 1:500 | #10326, IBL |
| EEA1 | 1:100 | #3288, Cell Signaling |
| Iba1 | 1:200 | #013-27691, Wako |
| GFAP | 1:200 | #MAB3402, Millipore |
| PSD-95 | 1:50 | #124011, Synaptic Systems |
| synaptophysin | 1:200 | #61012, PROGEN |

## REFERENCES

1. Alzheimer, A., Stelzmann, R.A., Schnitzlein, H.N. & Murtagh, F.R. An English translation of Alzheimer’s 1907 paper, “Uber eine eigenartige Erkankung der Hirnrinde”. Clin Anat 8, 429–431 (1995).

2. Sevigny, J., et al. The antibody aducanumab reduces Abeta plaques in Alzheimer’s disease. Nature 537, 50–56 (2016).

3. Sasaguri, H., et al. APP mouse models for Alzheimer’s disease preclinical studies. Embo j 36, 2473–2487 (2017).

4. Saito, T., et al. Single App knock-in mouse models of Alzheimer’s disease. Nat Neurosci 17, 661–663 (2014).

5. Li, S., et al. Swedish mutant APP-based BACE1 binding site peptide reduces APP β-cleavage and cerebral Aβ levels in Alzheimer’s mice. Sci Rep 5, 11322 (2015).

6. Kwart, D., et al. A Large Panel of Isogenic APP and PSEN1 Mutant Human iPSC Neurons Reveals Shared Endosomal Abnormalities Mediated by APP β-CTFs, Not Aβ. Neuron 104, 256–270.e255 (2019).

7. Lauritzen, I., et al. Intraneuronal aggregation of the β-CTF fragment of APP (C99) induces Aβ- independent lysosomal-autophagic pathology. Acta Neuropathol 132, 257–276 (2016).

8. Cataldo, A.M., et al. Endocytic pathway abnormalities precede amyloid beta deposition in sporadic Alzheimer’s disease and Down syndrome: differential effects of APOE genotype and presenilin mutations. Am J Pathol 157, 277–286 (2000).

9. Rogaeva, E., et al. The neuronal sortilin-related receptor SORL1 is genetically associated with Alzheimer disease. Nat Genet 39, 168–177 (2007).

10. Israel, M.A., et al. Probing sporadic and familial Alzheimer’s disease using induced pluripotent stem cells. Nature 482, 216–220 (2012).

11. Small, S.A., Simoes-Spassov, S., Mayeux, R. & Petsko, G.A. Endosomal Traffic Jams Represent a Pathogenic Hub and Therapeutic Target in Alzheimer’s Disease. Trends Neurosci 40, 592–602 (2017).

12. Toh, W.H. & Gleeson, P.A. Dysregulation of intracellular trafficking and endosomal sorting in Alzheimer’s disease: controversies and unanswered questions. Biochem J 473, 1977–1993 (2016).

13. Garneau, J.E., et al. The CRISPR/Cas bacterial immune system cleaves bacteriophage and plasmid DNA. Nature 468, 67–71 (2010).

14. Komor, A.C., Badran, A.H. & Liu, D.R. CRISPR-Based Technologies for the Manipulation of Eukaryotic Genomes. Cell 169, 559 (2017).

15. Scott, J.D., et al. Discovery of the 3-Imino-1,2,4-thiadiazinane 1,1-Dioxide Derivative Verubecestat (MK-8931)-A β-Site Amyloid Precursor Protein Cleaving Enzyme 1 Inhibitor for the Treatment of Alzheimer’s Disease. J Med Chem 59, 10435–10450 (2016).

16. Kennedy, M.E., et al. The BACE1 inhibitor verubecestat (MK-8931) reduces CNS beta-amyloid in animal models and in Alzheimer’s disease patients. Sci Transl Med 8, 363ra150 (2016).

17. Cradick, T.J., Qiu, P., Lee, C.M., Fine, E.J. & Bao, G. COSMID: A Web-based Tool for Identifying and Validating CRISPR/Cas Off-target Sites. Mol Ther Nucleic Acids 3, e214 (2014).

18. Bae, S., Park, J. & Kim, J.S. Cas-OFFinder: a fast and versatile algorithm that searches for potential off-target sites of Cas9 RNA-guided endonucleases. Bioinformatics 30, 1473–1475 (2014).

19. Yamakawa, H., Yagishita, S., Futai, E. & Ishiura, S. beta-Secretase inhibitor potency is decreased by aberrant beta-cleavage location of the “Swedish mutant” amyloid precursor protein. J Biol Chem 285, 1634–1642 (2010).

20. Hussain, I., et al. Oral administration of a potent and selective non-peptidic BACE-1 inhibitor decreases beta-cleavage of amyloid precursor protein and amyloid-beta production in vivo. J Neurochem 100, 802–809 (2007).

21. Rabe, S., et al. The Swedish APP mutation alters the effect of genetically reduced BACE1 expression on the APP processing. J Neurochem 119, 231–239 (2011).

22. Kalimo, H., et al. The Arctic AbetaPP mutation leads to Alzheimer’s disease pathology with highly variable topographic deposition of differentially truncated Abeta. Acta Neuropathol Commun 1, 60 (2013).

23. Hung, C.O.Y. & Livesey, F.J. Altered γ-Secretase Processing of APP Disrupts Lysosome and Autophagosome Function in Monogenic Alzheimer’s Disease. Cell Rep 25, 3647–3660.e3642 (2018).

24. Jiang, Y., et al. Alzheimer’s-related endosome dysfunction in Down syndrome is Abeta-independent but requires APP and is reversed by BACE-1 inhibition. Proc Natl Acad Sci U S A 107, 1630–1635 (2010).

25. Kim, S., et al. Evidence that the rab5 effector APPL1 mediates APP-βCTF-induced dysfunction of endosomes in Down syndrome and Alzheimer’s disease. Mol Psychiatry 21, 707–716 (2016).

26. Woodruff, G., et al. Defective Transcytosis of APP and Lipoproteins in Human iPSC-Derived Neurons with Familial Alzheimer’s Disease Mutations. Cell Rep 17, 759–773 (2016).

27. Xu, W., et al. Amyloid precursor protein-mediated endocytic pathway disruption induces axonal dysfunction and neurodegeneration. J Clin Invest 126, 1815–1833 (2016).

28. Treusch, S., et al. Functional links between Aβ toxicity, endocytic trafficking, and Alzheimer’s disease risk factors in yeast. Science 334, 1241–1245 (2011).

29. Willen, K., et al. Abeta accumulation causes MVB enlargement and is modelled by dominant negative VPS4A. Mol Neurodegener 12, 61 (2017).

30. Marshall, K.E., Vadukul, D.M., Staras, K. & Serpell, L.C. Misfolded amyloid-β-42 impairs the endosomal-lysosomal pathway. Cell Mol Life Sci (2020).

31. Baglietto-Vargas, D., et al. Generation of a humanized Abeta expressing mouse demonstrating aspects of Alzheimer’s disease-like pathology. Nat Commun 12, 2421 (2021).

32. Knupp, A., et al. Depletion of the AD Risk Gene SORL1 Selectively Impairs Neuronal Endosomal Traffic Independent of Amyloidogenic APP Processing. Cell Rep 31, 107719 (2020).

33. Pensalfini, A., et al. Endosomal Dysfunction Induced by Directly Overactivating Rab5 Recapitulates Prodromal and Neurodegenerative Features of Alzheimer’s Disease. Cell Rep 33, 108420 (2020).

34. Jonsson, T., et al. A mutation in APP protects against Alzheimer’s disease and age-related cognitive decline. Nature 488, 96–99 (2012).

35. Elvang, A.B., et al. Differential effects of gamma-secretase and BACE1 inhibition on brain Abeta levels in vitro and in vivo. J Neurochem 110, 1377–1387 (2009).

36. Hsu, P.D., et al. DNA targeting specificity of RNA-guided Cas9 nucleases. Nat Biotechnol 31, 827–832 (2013).

37. Yang, H., Wang, H. & Jaenisch, R. Generating genetically modified mice using CRISPR/Cas-mediated genome engineering. Nat Protoc 9, 1956–1968 (2014).

